# Using polyacrylamide hydrogels to model physiological aortic stiffness reveals that microtubules are critical regulators of isolated smooth muscle cell morphology and contractility

**DOI:** 10.1101/2021.12.14.472278

**Authors:** Sultan Ahmed, Robert. T. Johnson, Reesha Solanki, Teclino Afewerki, Finn Wostear, Derek T. Warren

**Author notes:** Corresponding author: Dr. Robert Johnson, School of Pharmacy, University of East Anglia, Norwich Research Park, UK. NR4 7TJ. Contributed equally to this work (Sultan Ahmed, Robert. T. Johnson).

## Abstract

Vascular smooth muscle cells (VSMCs) are the predominant cell type in the medial layer of the aortic wall and normally exist in a quiescent, contractile phenotype where actomyosin-derived contractile forces maintain vascular tone. However, VSMCs are not terminally differentiated and can dedifferentiate into a proliferative, synthetic phenotype. Actomyosin force generation is essential for the function of both phenotypes. Whilst much is already known about the mechanisms of VSMC actomyosin force generation, existing assays are either low throughput and time consuming, or qualitative and inconsistent. In this study, we use polyacrylamide hydrogels, tuned to mimic the physiological stiffness of the aortic wall, in a VSMC contractility assay. Isolated VSMC area decreases following stimulation with the contractile agonists angiotensin II or carbachol. Importantly, the angiotensin II induced reduction in cell area correlated with increased traction stress generation. Inhibition of actomyosin activity using blebbistatin or Y-27632 prevented angiotensin II mediated changes in VSMC morphology, suggesting that changes in VSMC morphology and actomyosin activity are core components of the contractile response. Furthermore, we show that microtubule stability is an essential regulator of isolated VSMC contractility. Treatment with either colchicine or paclitaxel uncoupled the morphological and/or traction stress responses of angiotensin II stimulated VSMCs. Our findings support the tensegrity model and we demonstrate that microtubules act to balance the actomyosin-derived traction stress generation and regulate the morphological responses of VSMCs.

## Introduction

Vascular smooth muscle cells (VSMCs) are the predominant cell type in the medial layer of the arterial wall. VSMCs normally exist in a quiescent, contractile phenotype where actomyosin-derived contractile forces maintain vascular tone (Brozovich et al., 2016). However, VSMCs are not terminally differentiated and can down-regulate contractile markers and dedifferentiate into a proliferative, synthetic phenotype (Rzucidlo et al., 2007; Liu et al., 2015; Shi and Chen, 2016). Both phenotypes retain the ability to generate actomyosin force that is essential for both VSMC contraction and migration (Ahmed and Warren, 2018; Afewerki et al., 2019). Contractile VSMCs possess a greater abundance of α-smooth muscle actin (αSMA) and smooth muscle-myosin heavy chain (SM-MyHC), that enhance their ability to generate actomyosin forces and contract (Rensen et al., 2007). However, both contractile and proliferative VSMCs generate actomyosin force via stimulating interactions between myosin II and filamentous actin. This process is regulated in both phenotypes by blood-borne factors, such as angiotensin II, that bind to receptors on the VSMC surface and mechanical factors including matrix stiffness and circumferential tension of the aortic wall (Qiu et al., 2010; Brozovich et al., 2016; Ahmed and Warren, 2018).

Young’s modulus is a measure of material stiffness. Atomic force microscopy studies have determined that the Young’s modulus of the medial layer of an elastic artery is between 5-37 kPa (Tracqui et al., 2011; Bae et al., 2016; Rezvani-Sharif et al., 2019). Microenvironment rigidity transmits ‘outside-in’ resistive forces to VSMCs and this process is dependent on focal adhesions that convey force between the ECM and cytoskeleton (Lacolley et al., 2017; Ahmed and Warren, 2018; Mohammed et al., 2019). VSMCs and other cell types respond to outside-in signals by exerting actomyosin based contractile forces on the matrix (inside-out forces) that scale with the outside-in resistive forces (Sun et al., 2008; Holle et al., 2018). Matrix rigidity rapidly activates Rho/ROCK signalling at ECM adhesions, initiating actin polymerisation and myosin light chain phosphorylation, thereby augmenting actomyosin activity (Ahmed and Warren, 2018). Actin cytoskeletal reorganisation and enhanced actomyosin activity increase VSMC integrated traction forces, the force VSMCs apply to the ECM (Li et al., 2020; Johnson et al., 2021). While much is known about VSMC actomyosin and contractile responses, we still lack an understanding of how VSMC structural and signalling components integrate to regulate these processes.

These actomyosin derived contractile forces place stress and intracellular tension upon the cell. In other cell types, these deformational forces are proposed to be balanced by the ECM and the microtubule network (Johnson et al., 2021). This relationship is described by the tensegrity model, whereby microtubules act as compression bearing struts, capable of resisting strain generated by the actin cytoskeleton (Stamenović, 2005; Brangwynne et al., 2006). The balance between compression and strain defines cell shape and stability (Stamenović, 2005). This model predicts that microtubule destabilisation will increase actomyosin derived force generation. In support of this, treatment with microtubule destabilisers, such as colchicine, result in enhanced force generation within coronary and aortic vessels (Sheridan et al., 1996; Platts et al., 1999, 2002; Paul et al., 2000; Zhang et al., 2000). However, our understanding of the role of microtubules in VSMC contractile agonist responses remains limited and some data contradicts the tensegrity model. For example, wire myography showed that microtubule stabilisation had no effect on the ability of isolated aortic rings to contract (Zhang et al., 2000). Dynamic instability, the ability of microtubules to constantly cycle through phases of growth and shrinkage, is an inherent characteristic that enables the microtubule network to rapidly reorganise in response to the changing mechanical requirements of the cell (Nogales, 2001). Whilst observations regarding the function of microtubule stability in VSMC contraction have been made, our understanding of the mechanisms behind these observations remains incomplete.

One of the largest obstacles for identifying different mechanisms of VSMC contraction remains the lack of *in vitro* tools. The current gold standard for assessing isolated cell actomyosin activity is traction force microscopy (TFM) (Muhamed et al., 2017). TFM is used to quantitatively measure the stress exerted by a cell on its substrate, which is then used as an indicator of cell contractility (Kraning-Rush et al., 2012; Lekka et al., 2021). Additionally, atomic force microscopy (AFM) has been used to assess actomyosin cortical tension and the force transduced through individual FA complexes (Sanyour et al., 2019). However, these techniques have several limitations, mainly being time consuming and low throughput (Haase and Pelling, 2015; Colin-York and Fritzsche, 2018; Schierbaum et al., 2019). Collagen gel assays, that are easily performed with generic lab equipment and skills, provide an alternative to these techniques. Typical collagen assays involve the suspension of a cell population in a prefabricated collagen gel. Contraction is then assessed by observing dimensional changes of the gel (Ngo et al., 2006). Limitations of collagen gels primarily relate to the qualitative nature of the assay and inconsistencies in gel shape. Collagen gels also lack rigidity control and are softer than the physiological arterial wall. These limitations severely affect the reproducibility and reliability of such assays (Vernon and Gooden, 2002).

Previous studies have reported that isolated VSMCs display a reduction in cell area upon contractile agonist stimulation (Li et al., 1999; Wang et al., 2017; Halaidych et al., 2019). Suggesting that changes in VSMC morphology and actomyosin activity are important components of the VSMC contractile response. Given their limitations, existing *in vitro* VSMC contractility assays are not able to investigate these processes. In this study we develop and validate an approach for screening isolated VSMC contraction using cell area as a reporter. The assay uses polyacrylamide hydrogels that are easily fabricated and mimic the rigidity of the physiological elastic arterial wall (Minaisah et al., 2016; Porter et al., 2020). Moreover, we show that combined with TFM, this technique allows investigation into factors that regulate both VSMC morphology and traction stress generation. Finally, we identify microtubule stability as a key regulator of VSMC contractile responses.

## Materials and Methods

### Polyacrylamide hydrogel preparation

Hydrogels were prepared as described previously (Minaisah et al., 2016). Briefly 30 mm coverslips were activated by treating with (3-Aminopropyl)triethoxysilane for 2 minutes, washed 3x in dH_2_O and fixed in 0.5% glutaraldehyde for 40 minutes. Coverslips were subsequently washed 3x in dH_2_O and left to air dry. The required volume of 12 kPa polyacrylamide hydrogel mix (7.5% acrylamide, 0.15% bis-acrylamide in water) was supplemented with 10% APS (1:100) and TEMED (1:1000) before 50 µl of the solution was placed on a standard microscopy slide and covered by an activated coverslip. Once set, the fabricated 12 kPa hydrogel was removed, placed into a 6-well plate, washed 3x with dH_2_O and crosslinked with sulfo-SANPAH (1:3000) under UV illumination (365 nm). Finally, hydrogels were washed with PBS and functionalised with collagen I (0.1 mg/ml) for 10 minutes at room temperature. Hydrogel stiffness was previously confirmed using a JPK Nanowizard-3 atomic force microscope (Porter et al., 2020).

### Vascular Smooth Muscle Cell culture and drug treatments

Human adult aortic VSMCs (passage 3-10) were purchased from Cell Applications Inc (354-05a). VSMCs were grown in growth media (Cell Applications Inc), prior to being washed with Earle’s Balanced Salt Solution (Thermo) and seeded in basal media (Cell Applications Inc) onto 12 kPa hydrogels, 18 hours prior to drug treatment. Standard VSMCs culture was performed as described previously (Ragnauth et al., 2010; Warren et al., 2015).

Quiescent VSMCs were stimulated with either angiotensin II (0.01 µM – 100 µM) (Merck) or carbachol (0.01 µM – 100 µM) (Merck) for 30 minutes. For all other drug treatments, quiescent VSMCs were pretreated with the stated dose for 30 minutes, prior to co-treatment with angiotensin II (10 µM) for an additional 30 minutes. Please see Supplementary Table 1 for a list of compounds used in this study.

### Immunofluorescence and VSMC Area/Volume Analysis

Following treatment, cells were fixed in 4% paraformaldehyde (actin cytoskeleton) or ice-cold methanol (microtubules), permeabilised with 0.5% NP40 and blocked in 3% BSA/PBS. Targets were visualized using antibodies raised against lamin A/C (SAB4200236, Sigma), or α-tubulin (3873S, CST) in combination with a relevant Alexa Fluor 488 secondary antibody (A1101 or A1103, Thermo). F-actin was stained using Rhodamine Phalloidin (R145, Thermo). All images were captured at 20x (cell area/volume) or 40x (microtubule organization) magnification using either a Zeiss LSM510-META or a Zeiss LSM980-Airyscan confocal microscope. Cell area was measured using FIJI, open-source software (Schindelin et al., 2012), whilst cell volume was calculated using Volocity 6.3.

### Cold-stable Microtubule Stability Assay

The number of cold-stable microtubules per cell was determined as previous (Atkinson et al., 2018). Briefly, after treatment, cells were placed on ice for 15 minutes before being washed once with PBS and twice with PEM buffer (80 µM PIPES pH 6.8, 1 mM EGTA, 1 mM MgCl_2_, 0.5% Triton X-100 and 25% (w/v) glycerol) for 3 minutes. Cells were then fixed for 20 minutes in ice-cold MeOH and prepared for staining as detailed above. Cell nuclei were identified using DAPI. Images were captured at 40x magnification using an Axioplan Epifluorescent microscope.

### Traction Force Microscopy

For Traction Force Microscopy (TFM), VSMCs were seeded onto 12 kPa hydrogels containing 0.5 µm red fluorescent (580/605) FluoSpheres (1:1000) (Invitrogen). Imaging was performed using a Zeiss Axio Observer live cell imaging system that captured 20x magnification images every 2 minutes. TFM was performed either in real-time or as an end point assay whereby images were captured before and after cell lysis. Lysis was achieved by the addition of 0.5% Triton X-100. Drift was corrected using the ImageJ StackReg plugin and traction force was calculated using the ImageJ plugin described previously to measure FluoSphere displacement (Tseng et al., 2012). Briefly, bead displacement was measured using the first and last image of the movie sequence. The cell region was segmented by overlaying the traction map with the cell image, highlighting the cell traction region with an ROI and extracting the traction forces in each pixel by using the save XY coordinate function in ImageJ (Porter et al., 2020).

### Cell viability assay

Cell viability was determined using a RealTime-Glo™ MT Cell Viability Assay following manufacturers instruction. Briefly, 5,000 cells per well in a 96-well plate were exposed to a range of drug concentrations for 1 hr, prior to luminescence being measured using a Wallac EnVision 2103 Multilabel Reader (PerkinElmer).

### Statistics

All statistical analyses were performed in GraphPad Prism 6.05. Dose response curves are presented as the mean data +/-SEM (error bars) plotted on a logarithmic dose response scale. EC_50_ and IC_50_ data were generated by non-linear regression. Results are presented as mean +/-SEM. Dot plot graphs are presented to show data distribution, with each point corresponding to an individual cell measurement. Bars indicate the mean value, +/-SEM (error bars). Comparison of multiple groups was achieved using a one-way ANOVA analysis with Bonferroni post-hoc test. For comparison of two groups, unpaired student t-tests were performed. Analysis of the real time Angiotensin II contraction data was performed using non-linear regression. A one phase association analysis was performed on traction stress vs time data and a one phase decay analysis was performed on the VSMC area vs time data.

## Results

### Contractile agonist stimulation reduces VSMC area on pliable hydrogels

We set out to develop a screen, performed at a physiologically relevant rigidity, to unravel mechanisms regulating VSMC actomyosin activity and contraction (**Supplementary Figure S1**). Quiescent VSMCs grown on pliable, 12 kPa hydrogels were treated with a concentration range of the contractile agonists, angiotensin II or carbachol. Contractile response was measured through changes in VSMC area, which previous studies have shown correlates with isolated VSMC contractile activity (Li et al., 1999; Wang et al., 2017; Halaidych et al., 2019). Analysis confirmed that contractile agonist stimulation resulted in a reduction in VSMC area (**Figures 1A, B and Supplementary Figures S2A and B**). Further analysis of VSMC area revealed the assay could be used to determine the EC_50_ values of both angiotensin II and carbachol (**Figure 1C and Supplementary Figure S2C**), confirming the assay could be used to measure and compare agonist potency. Subsequent experiments were performed with 10 µM angiotensin II, a dose that produced the maximal response (**Figures 1A and B**). To observe the effect of angiotensin II stimulation on isolated VSMC volume, we next performed confocal microscopy. Analysis revealed that angiotensin II stimulated VSMCs displayed a reduction in area, but no change in volume (**Supplementary Figure S3**).

**Figure 1:**
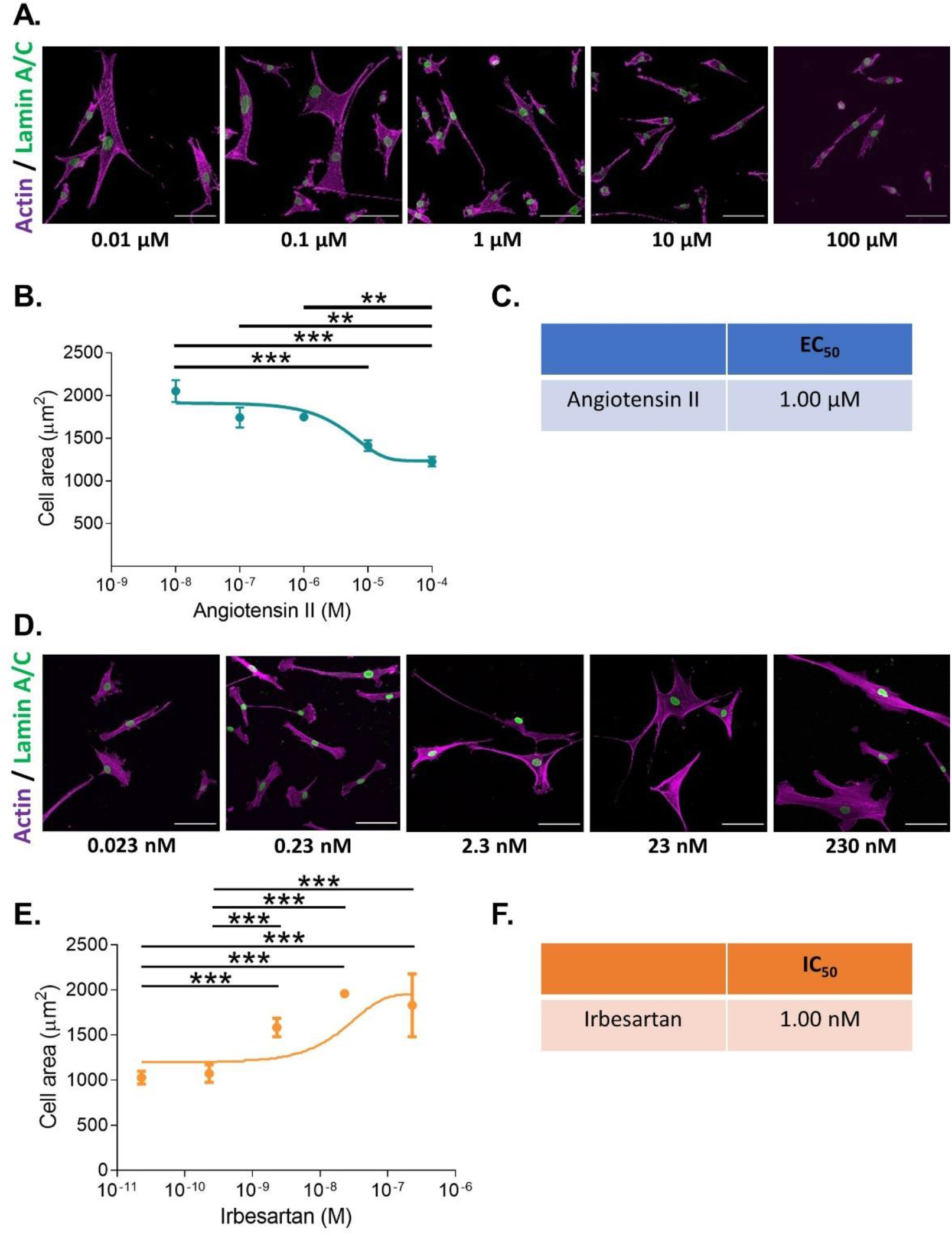
VSMCs grown on pliable hydrogels display decreased area upon stimulation with the contractile agonist angiotensin II. **(A)** Representative images of isolated VSMCs cultured on 12 kPa polyacrylamide hydrogels treated with a range of angiotensin II concentrations for 30 minutes. Actin cytoskeleton (purple) and Lamin A/C labelled nuclei (green). Scale bar = 100 μm. **(B)** Isolated VSMC area, representative of 3 independent experiments with ≥ 150 cells analysed per condition. **(C)** EC_50_ of angiotensin II calculated from **B. (D)** Representative images of isolated VSMCs cultured on 12 kPa hydrogels and treated with angiotensin II (10 µM) for 30 minutes in the presence of a range of irbesartan concentrations. Actin cytoskeleton (purple) and Lamin A/C (green). Scale bar = 100 μm. **(E)** Isolated VSMC area, representative of 3 independent experiments with ≥ 50 cells analysed per condition. **(F)** IC_50_ of irbesartan calculated from **E**. (** = *p*<0.01), (*** = *p*<0.001).

To confirm that the change in area was specific for receptor activation, we utilised the angiotensin II antagonist irbesartan and the cholinergic antagonist, atropine. VSMCs grown on pliable hydrogels were stimulated with angiotensin II or carbachol in the presence of an increasing dose of irbesartan or atropine, respectively. As expected, irbesartan and atropine treatment prevented a reduction in VSMC area, confirming that these changes were driven by receptor activation (**Figures 1D-F and Supplementary Figure S4**).

### Myosin II mediated traction stress drives changes in Angiotensin II stimulated VSMC area

The above data demonstrated the validity of our approach in generating *EC*_*50*_ and *IC*_*50*_ data for agonists/antagonists of isolated VSMC contractile function. We next sought to confirm that changes in VSMC area were due to contraction and not due to membrane retraction. To do this, we performed traction force microscopy to measure the displacement of beads embedded within the hydrogels. Analysis revealed that angiotensin II treatment stimulated VSMCs to contract, marked by a reduction in area (**Figures 2A and B**). Minimum cell area was observed 8 minutes after treatment (**Figures 2A and B**). Importantly, beads moved towards VSMCs and both maximal and integrated traction stress increased rapidly and plateaued after 8 minutes (**Figures 2A, C and D**). This data confirmed the correlation between reduced VSMC area and traction stress generation upon angiotensin II stimulation. To further confirm that actomyosin derived force was driving these changes in VSMC area, we next used the myosin II inhibitor blebbistatin and the ROCK inhibitor Y-27632. As expected, treatment with either blebbistatin or Y-27632 blocked the angiotensin II mediated reduction in area, however, actomyosin inhibited VSMCs possessed an increased volume, compared to their angiotensin II treated counterparts (**Figure 3**). This confirmed that actomyosin activation was driving VSMC contraction in our system.

**Figure 2:**
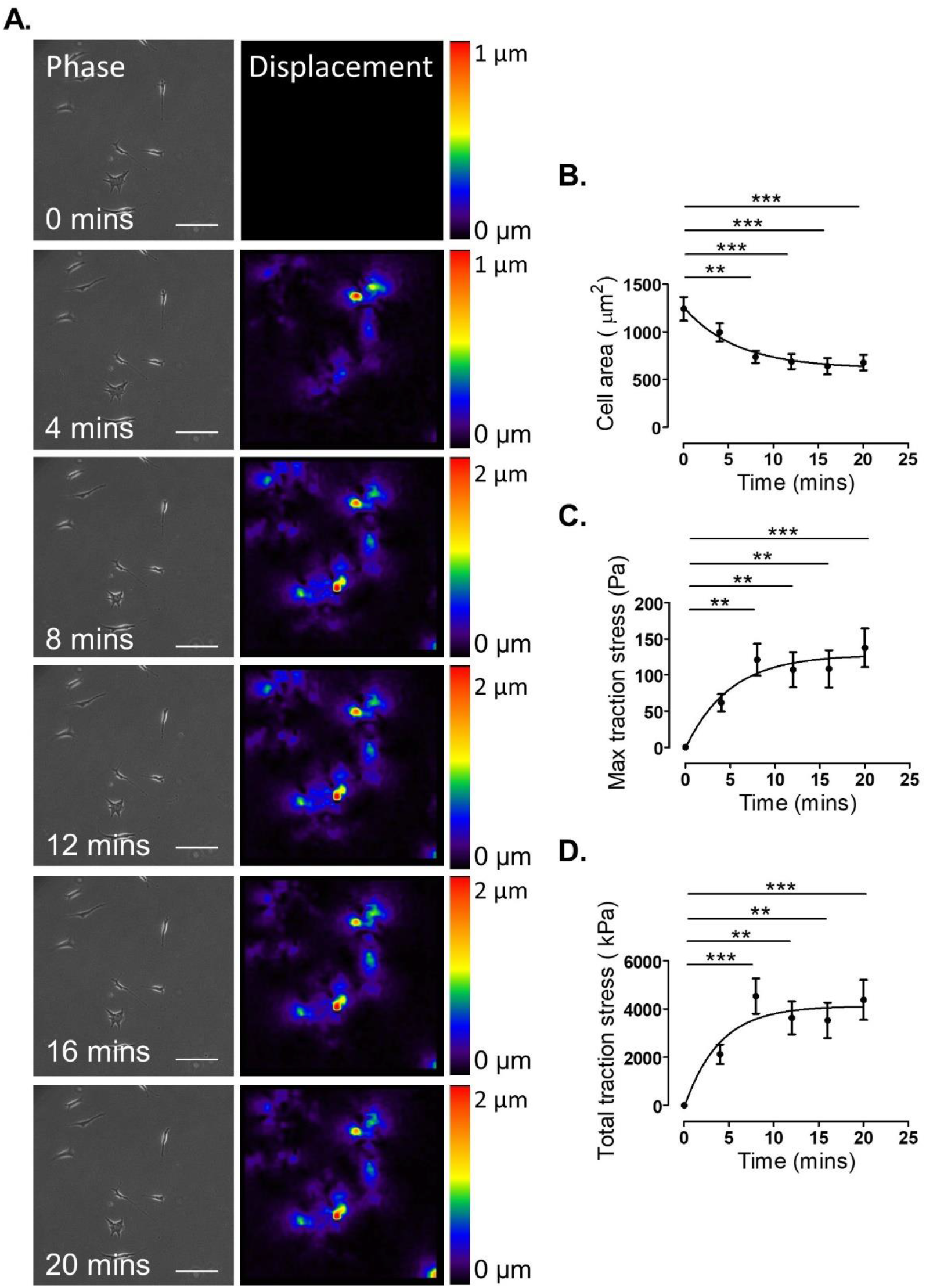
Traction stress generation correlates with decreased VSMC area following angiotensin II stimulation. **(A)** Representative phase images and bead displacement heat maps of isolated VSMCs grown on 12 kPa polyacrylamide hydrogels and stimulated with angiotensin II over a 20 minute time course. Scale bar = 100 μm. Displacement heat maps were generated by comparing the bead displacement for each time point to their initial positions at t=0, as such there is no displacement at t=0. **(B)** Isolated VSMC area, **(C)** maximum traction stress and **(D)** total traction stress generated at each time point compared to t=0. Graphs represent the combined data from 3 independent experiments, with measurements taken from 35 VSMCs. (** = *p*<0.01), (*** = *p*<0.001).

**Figure 3.**
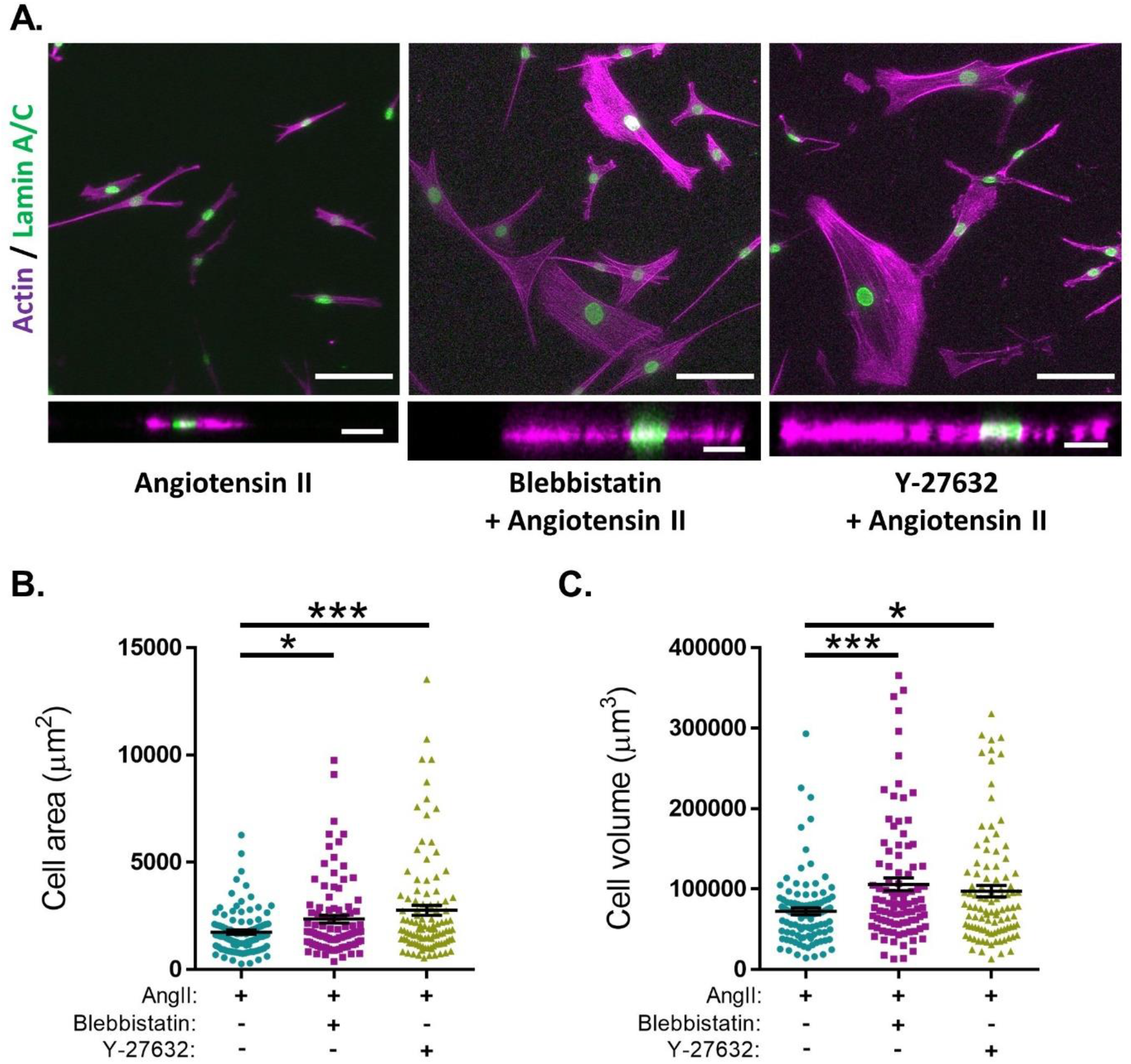
Actomyosin inhibition prevents angiotensin II induced changes in VSMC area and increases VSMC volume on pliable hydrogels. **(A)** Representative images of isolated VSMCs cultured on 12 kPa polyacrylamide hydrogels pre-treated with either blebbistatin or Y-27632 for 30 minutes prior to cotreatment with angiotensin II (AngII) (10 µM) for an additional 30 minutes. Actin cytoskeleton (Rhodamine phalloidin, Purple) and Lamin A/C (green). **Top -** Representative XY images of VSMC area. Scale bar = 100 μm. **Bottom -** Representative XZ images of VSMC height. Scale bar = 30 μm. **(B)** Isolated VSMC area and **(C)** Isolated VSMC volume. Both **B&C** are representative of 3 independent experiments with ≥ 95 cell analysed per condition. (* = *p*<0.05), (*** = *p*<0.001).

### Microtubule destabilisation alters the morphological response of angiotensin II stimulated VSMCs

Microtubule depolymerisation increases the constriction and myogenic tone of aortic and carotid tissues (Leite and Webb, 1998; Platts et al., 2002; Johnson et al., 2021). However, the precise impact of microtubule disruption on VSMC contraction remains unknown. To address this, we determined the impact of angiotensin II stimulated VSMC contraction upon microtubule organisation and stability. Angiotensin II stimulation promoted the microtubule network to reorganise, switching from a straight, elongated arrangement of parallel microtubules to a more interlinking meshwork of microtubules (**Supplementary Figure S5)**. Although angiotensin II stimulation induced the reorganisation of the microtubule cytoskeleton, there was no change in the number of cold-stable microtubules between quiescent and angiotensin II stimulated VSMCs (**Figures 4A and B**). Next, we used our assay to test the impact of microtubule disruption on VSMC contraction. Colchicine, a microtubule destabiliser, induced a concentration dependent increase in angiotensin II treated VSMC area, suggesting that microtubule depolymerisation was promoting VSMC relaxation (**Figures 4C-E**). In contrast, paclitaxel, a microtubule stabiliser, had no effect on the area of angiotensin II treated VSMCs (**Figures 4F and G**). Analysis confirmed that the concentrations of colchicine or paclitaxel used were sufficient to reduce or increase the number of cold-stable microtubules respectively (**Supplementary Figure S6A-D**). Furthermore, these concentrations produced no significant reductions in cell viability (**Supplementary Figure S6E & F**). Surprisingly, confocal microscopy revealed that colchicine increased VSMC volume (**Figures 5A-C**), whereas the volume of paclitaxel treated VSMCs remained unchanged (**Figures 5 D-F**).

**Figure 4.**
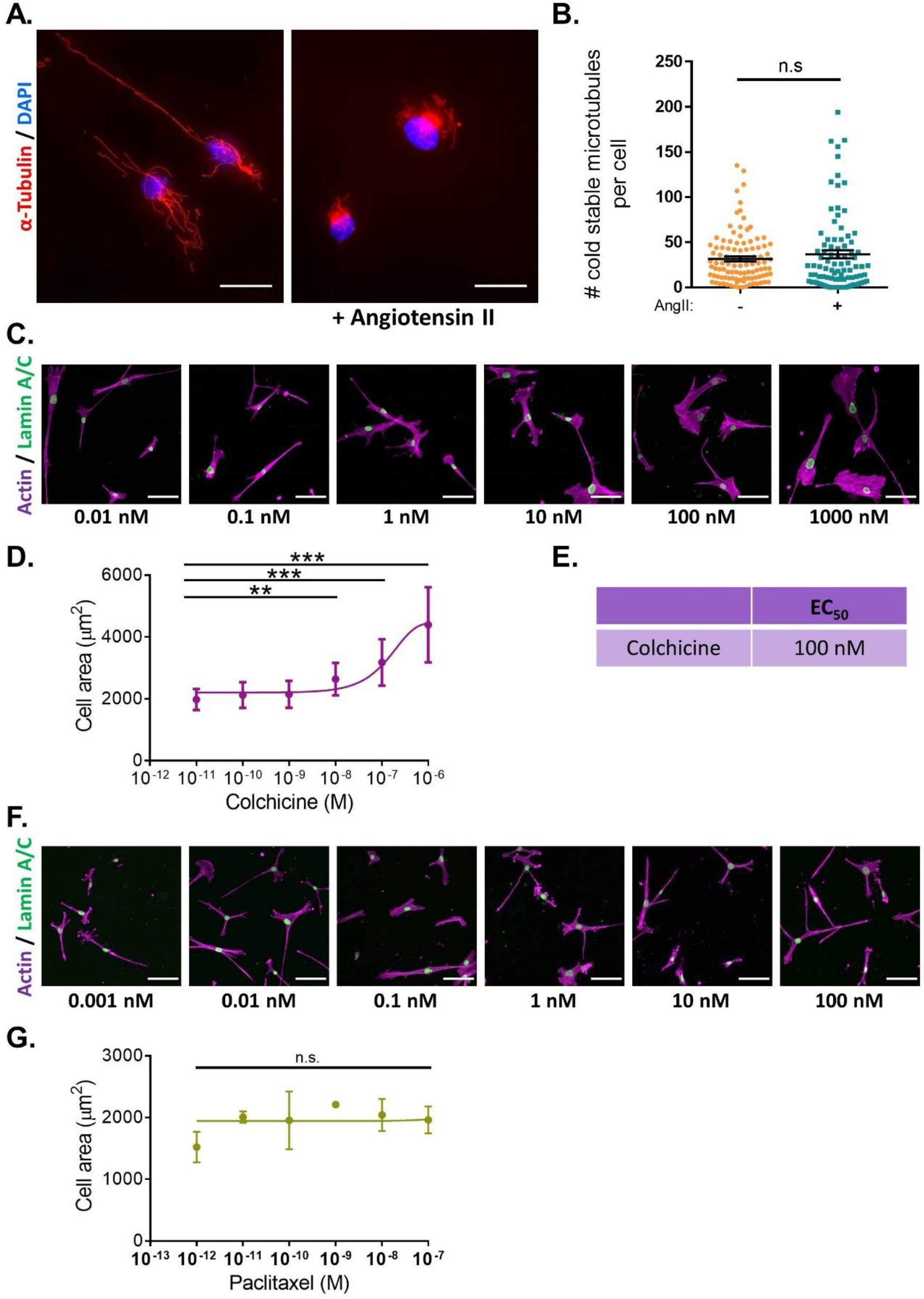
Microtubule destabilisation prevents angiotensin II induced changes in VSMC area. **(A)** Representative images of isolated VSMCs cultured on 12 kPa polyacrylamide hydrogels, stained for cold-stable microtubules (α-tubulin, red) and cell nuclei (DAPI, blue). Scale bar = 50 µm. Angiotensin II (AngII) (10 µM) stimulation was performed for 30 minutes. **(B)** Number of cold-stable microtubules per cell, representative of 4 independent experiments, with ≥ 90 cells analysed per condition. **(C)** Representative images of isolated VSMCs cultured on 12 kPa hydrogels pre-treated with a range of colchicine concentrations for 30 minutes prior to cotreatment with angiotensin II (10 µM) for an additional 30 minutes. Actin cytoskeleton (purple) and Lamin A/C (green). Scale bar = 100 μm. **(D)** Isolated VSMC area, representative of 3 independent experiments with ≥ 55 cells analysed per condition. **(E)** EC_50_ of colchicine calculated from **D. (F)** Representative images of isolated VSMCs cultured on 12 kPa hydrogels pre-treated with a range of paclitaxel concentrations for 30 minutes prior to cotreatment with angiotensin II (10 µM) for an additional 30 minutes. Actin cytoskeleton (purple) and Lamin A/C (green). Scale bar = 100 μm. **(G)** Isolated VSMC area, representative of 3 independent experiments with ≥ 65 cells analysed per condition. (n.s. = non-significant), (** = *p*<0.01), (*** = *p*<0.001).

**Figure 5.**
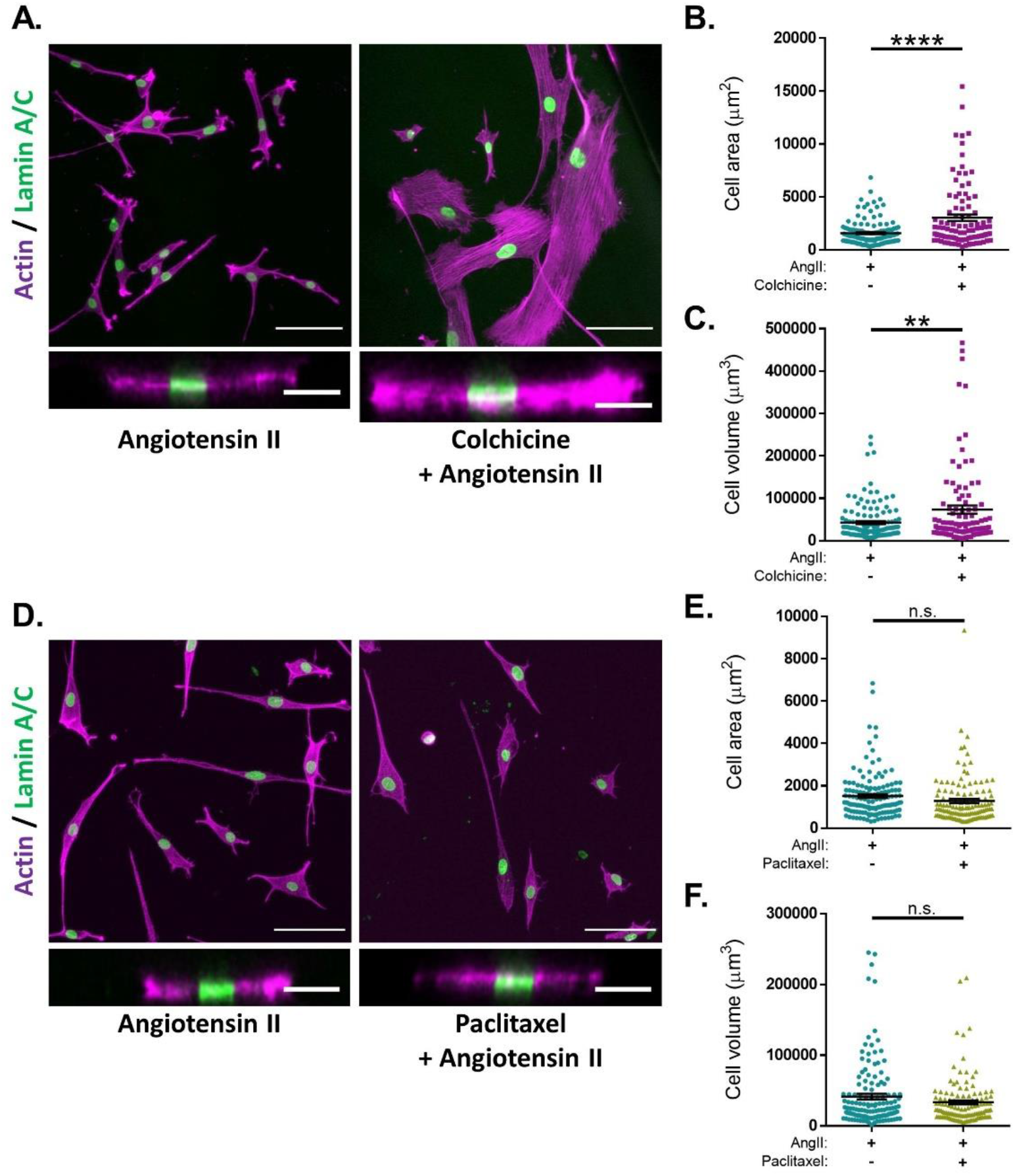
Microtubule destabilisation alters angiotensin II stimulated VSMC volume on pliable hydrogels. **(A)** Representative images of isolated VSMCs cultured on 12 kPa polyacrylamide hydrogels pre-treated +/-colchicine (100 nM) for 30 minutes prior to cotreatment with angiotensin II (AngII) (10 µM) for an additional 30 minutes. Actin cytoskeleton (purple) and Lamin A/C (green). **Top -** Representative XY images of VSMC area. Scale bar = 100 μm. **Bottom -** Representative XZ images of VSMC height. Scale bar = 30 μm. **(B)** Isolated VSMC area and **(C)** Isolated VSMC volume. Both **B&C** are representative of 3 independent experiments with ≥ 100 cell analysed per condition. **(D)** Representative images of isolated VSMCs cultured on 12 kPa polyacrylamide hydrogels pre-treated +/-paclitaxel (1 nM) for 30 minutes prior to cotreatment with angiotensin II (AngII) (10 µM) for an additional 30 minutes. Actin cytoskeleton (purple) and Lamin A/C (green). **Top -** Representative XY images of VSMC area. Scale bar = 100 μm. **Bottom -** Representative XZ images of VSMC height. Scale bar = 30 μm. **(E)** Isolated VSMC area and **(F)** Isolated VSMC volume. Both **E&F** are representative of 3 independent experiments with ≥ 125 cell analysed per condition. (n.s. = non-significant), (** = *p*<0.01), (**** = *p*<0.0001)

### Changes in microtubule stability alters the traction stress generation of isolated VSMC

The data above suggested that microtubule destabilisation was altering the angiotensin II induced area/volume response of isolated VSMCs. Previous studies using wire or pressure myography have reported that microtubule destabilisation increased VSMC force generation (Paul et al., 2000; Zhang et al., 2000; Platts et al., 2002). To confirm the impact of microtubule destabilisation on actomyosin activity, TFM was performed to measure traction stresses exerted by VSMCs. In agreement with previous studies, analysis revealed that pre-treatment with colchicine increased the maximal and total traction stress exerted by angiotensin II stimulated VSMCs (**Figure 6A-C**). Finally, we examined the effect of microtubule stabilisation on traction stress. Paclitaxel pre-treatment had no effect on maximal traction stress. However, total traction stress was reduced when compared to their untreated counterparts (**Figures 6D-F**). This confirmed that microtubule stability was essential for VSMC actomyosin activity and force generation.

**Figure 6.**
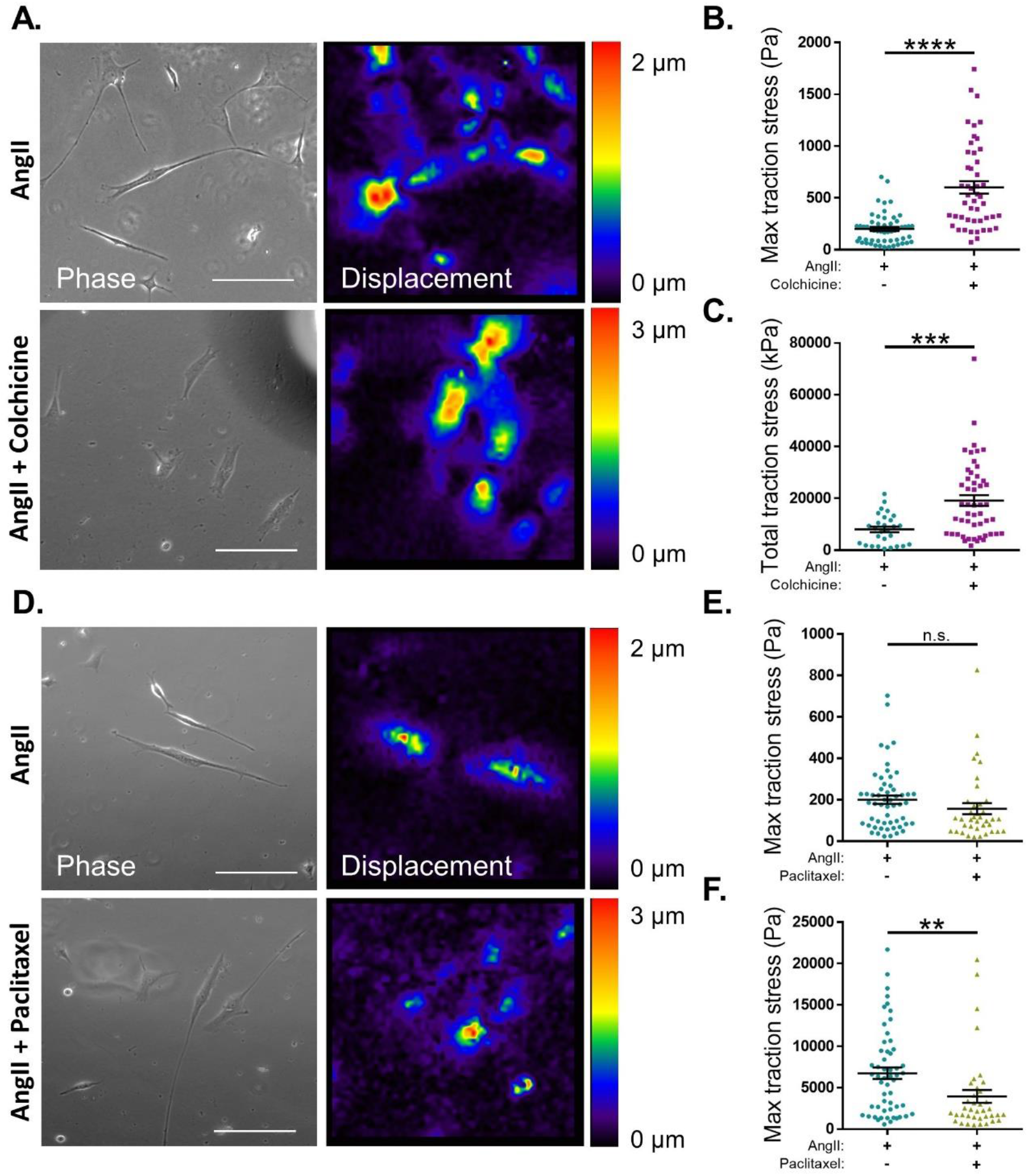
Microtubule stability regulates angiotensin II induced VSMC traction stress generation. **(A)** Representative phase images and bead displacement heat maps of isolated VSMCs grown on 12 kPa polyacrylamide hydrogels. Cells were pre-treated +/-colchicine (100 nM) for 30 minutes prior to cotreatment with angiotensin II (AngII) (10 µM) for an additional 30 minutes. Scale bar = 100 μm. **(B)** Maximum traction stress and **(C)** Total traction stress generation. Both **B&C** are representative of 3 independent experiments with ≥ 48 cell analysed per condition. **(D)** Representative phase images and bead displacement heat maps of isolated VSMCs grown on 12 kPa polyacrylamide hydrogels. Cells were pre-treated +/-paclitaxel (1 nM) for 30 minutes prior to cotreatment with angiotensin II (AngII) (10 µM) for an additional 30 minutes. Scale bar = 100 μm. **(E)** Maximum traction stress generation and **(F)** Total traction stress generation. Both **E&F** are representative of 3 independent experiments with ≥ 38 cell analysed per condition. (n.s. = non-significant), (** = *p*<0.01), (*** = *p*<0.001), (**** = *p*<0.0001).

## Discussion

Whilst much is known about VSMC contractile responses in physiology, *in vitro* assays to examine this process remain limited. Tissue culture plastic and glass are approximately a thousand times stiffer than the aortic wall (Minaisah et al., 2016), meaning that changes in VSMC area are driven by membrane retraction rather than contraction. Existing technologies to examine the contractile response of isolated VSMCs are either low throughput, such as TFM, or are inconsistent and lack rigidity control, such as the collagen gel assay (Vernon and Gooden, 2002; Colin-York and Fritzsche, 2018). We therefore set out to establish and validate a polyacrylamide hydrogel-based assay for screening the contractility of isolated VSMCs. Polyacrylamide hydrogels are widely used in cell biology and can be easily fabricated with generic research equipment and skills (Kandow et al., 2007; Caliari and Burdick, 2016; Minaisah et al., 2016; Mohammed et al., 2019). We demonstrate that changes in isolated VSMC area were driven by angiotensin II induced traction stress generation, that pulled the compliant hydrogels towards the VSMC, decreasing cell area **(Figures 1A, B, and 2)**. Demonstrating that, isolated VSMC area can be used as a reporter of contractility on pliable hydrogels. VSMC volume remained unchanged by contractile agonist stimulation **(Supplementary Figure S3)**. Importantly, this assay was sensitive enough to generate EC_50_ and IC_50_ information of contractile agonists and antagonists, respectively **(Figures 1C,F**, **S2C and S4C)**.

We have used this polyacrylamide hydrogel-based assay to investigate the role of microtubule stability in isolated VSMC morphology and traction stress generation. We identify microtubule stability as a critical regulator of isolated VSMC contractility and show that proper microtubule stability is essential to couple VSMC morphology and traction stress generation **(Figure 4A-D and 6A-C)**. Our findings fit the tensegrity model of cellular mechanics (Stamenović, 2005); microtubule destabilisation increased traction stress generation, whereas microtubule stabilisation had the opposite effect **(Figure 6)**. Our findings are supported by previous studies that show microtubule destabilising agents promoted an increase in isometric force generation by wire myography (Sheridan et al., 1996; Platts et al., 1999, 2002; Paul et al., 2000; Zhang et al., 2000). Whilst overall microtubule stability is maintained, angiotensin II induced contraction initiates remodelling of the microtubule network **(Supplementary Figure 5)**. Further work is required to understand the importance of this reorganisation, however, a recent study has demonstrated that stretched-induced reorganisation of VSMC microtubules enhances the efficiency of mitochondrial ATP production, thereby increasing actomyosin force generation (Bartolák-Suki et al., 2015).

However, contradictory to our findings **(Figure 6D-F)**, previous studies have shown that microtubule stabilisation does not alter VSMC force generation by wire myography (Zhang et al., 2000). In this study, we show that paclitaxel had no effect on VSMC morphology **(Figure 4F and G)** and this may potentially account for this discrepancy. Additionally, we identify the IC_50_ of colchicine to be within the nanomolar range (**Figure 4C-E)**. Paclitaxel also demonstrated significant microtubule stabilisation when used in the nanomolar range (**Supplementary Figure 6C and D)**. However, previous studies have used microtubule targeting agents in the tens of micromolar range (Platts et al., 1999; Paul et al., 2000; Zhang et al., 2000). Numerous studies have shown that these agents can be cytotoxic in micromolar and, in some cell lines, nanomolar ranges (Pasquier et al., 2006; Thomopoulou et al., 2016; Atkinson et al., 2018; Abbassi et al., 2019; Čermák et al., 2020). Meanwhile, in the present study we used these agents in nanomolar ranges, which had no effect on VSMC viability (**Supplementary Figure 6E & F)**. Surprisingly, wire myography has shown that VSMC death does not trigger a reduction in isometric force generation (Clarke et al., 2006). Previously, a tamoxifen model that induced VSMC death resulted in the loss of approximately two thirds of VSMCs. However, the remaining VSMCs generated more contractile forces, resulting in the maintenance of the overall isometric force generation (Clarke et al., 2006). This observation may account for the divergence from the tensegrity model predictions, as seen in previous studies (Zhang et al., 2000). Clearly more research is needed to clarify the role of the microtubules and cell death in regulating VSMC contractility and vascular tone.

As described above, our microtubule stabilisation/destabilisation traction stress data fits the tensegrity model of cellular mechanics **(Figure 6)**. However, we also show that microtubule destabilisation/stabilisation uncoupled the morphological response from the amount of traction stress VSMCs generated following angiotensin II stimulation **(Figure 5)**. In our assay, isolated VSMC volume was unchanged by contractile agonist stimulation and changes in area were driven by actomyosin induced contraction **(Figure 2 and S3)**. Microtubule destabilisation promoted increased VSMC volume **(Figure 5A and C)**, and we propose that this increased volume is enhancing spreading in these cells. We predicted that these cells were undergoing a hypertrophic response, however more work is needed to clarify how these cells respond over a longer time course. Surprisingly, inhibition of non-muscle myosin II or ROCK also resulted in enhanced VSMC volume **(Figure 3A and C)**. This suggests that actin and microtubule networks may mechanically cooperate to regulate VSMC volume control. Much is known about the synergistic nature of microtubules and actomyosin activity. Indeed, non-muscle myosin II has been shown supress microtubule growth to stabilise cell morphology (Sato et al., 2020), whilst microtubule acetylation has been found to regulate actomyosin-derived force generation (Joo and Yamada, 2014; Seetharaman et al., 2021). Actomyosin and microtubule cross-talk also regulates morphology during differentiation and cell migration in a variety of cell types (Akhshi et al., 2014; Wu and Bezanilla, 2018; Seetharaman and Etienne-Manneville, 2020), suggesting that communication between these mechanical networks is fundamentally important in determining VSMC morphology.

## Author Contributions

SA, RTJ, RS, TA, and FW were responsible for performing experiments and analysed data. DW analysed data. RTJ and DW were responsible for the design, writing and editing of this manuscript.

