## Supplementary Materials for "Using polyacrylamide hydrogels to model physiological aortic stiffness reveals that microtubules are critical regulators of isolated smooth muscle cell morphology and contractility"

### Supplementary Tables

#### Supplementary Table 1 – List of compounds used in this study

| Compound | Concentration Range | Optimal Concentration | Product Code | Supplier |
|---|---|---|---|---|
| Angiotensin II | 0.01 – 100 µM | 10 µM | A9525 | Merck |
| Atropine | 0.038 – 380 nM | - | ab145582 | Abcam |
| Blebbistatin | - | 40 µM | B0560 | Sigma |
| Carbachol | 0.01 – 100 µM | 10 µM | C4382 | Merck |
| Colchicine | 0.01 – 1000 nM | 100 nM | C9754 | Sigma |
| Irbesartan | 0.023 – 230 nM | - | I2286 | Merck |
| Paclitaxel | 0.001 – 100 nM | 1 nM | T7402 | Sigma |
| Y-27632 | - | 5 µM | Y0503 | Sigma |

**Supplementary Figures**

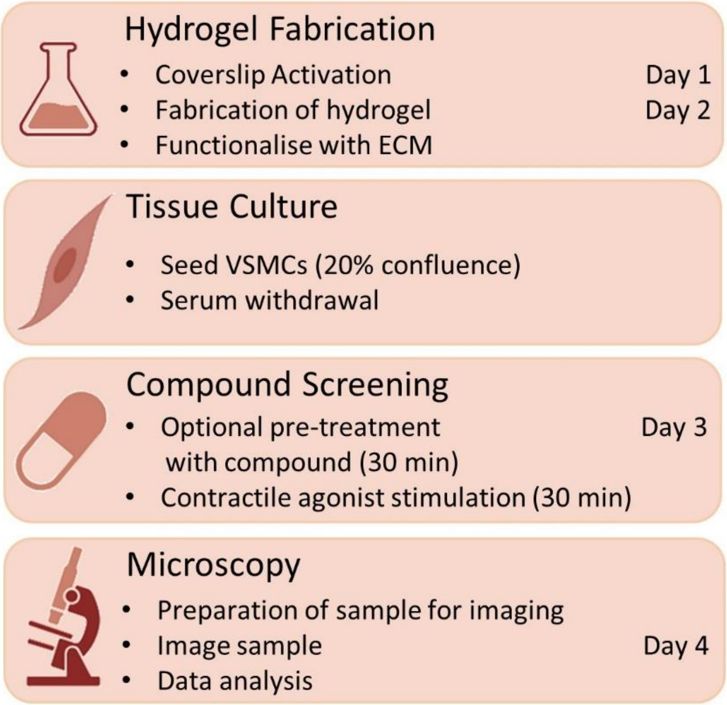


**Supplementary Figure S1: Workflow of the polyacrylamide hydrogel contractility assay.** VSMC contractility can be assayed using generic laboratory equipment and skills using polyacrylamide hydrogels. Briefly, glass coverslips are activated and air dried, enabling a hydrogel to be cast upon them. Hydrogels are then functionalised with an ECM component and seeded with VSMCs at low confluence. Serum withdrawal is performed overnight to induce quiescence and VSMCs are subsequently pre-treated with a compound of interest (optional) for 30 minutes, before cotreatment with a contractile agonist for an additional 30 minutes. Following treatment, cells are fixed and prepared for microscopic analysis.

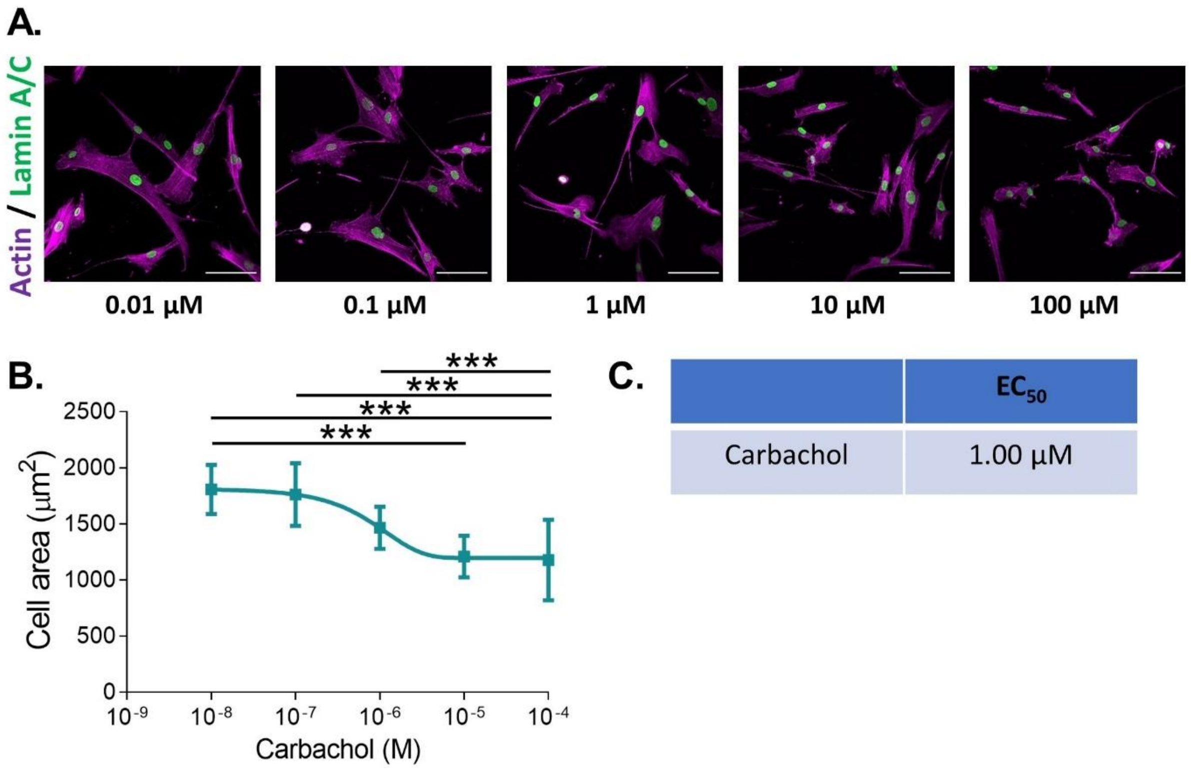


**Supplementary Figure S2: VSMCs grown on pliable hydrogels display decreased area upon stimulation with the contractile agonist carbachol. (A)** Representative images of isolated VSMCs cultured on 12 kPa, pliable polyacrylamide hydrogels and treated with a range of carbachol concentrations for 30 minutes. Actin cytoskeleton (Rhodamine phalloidin, Purple) and Lamin A/C (green). Scale bar = 100 µm. (**B)** Isolated VSMC area, representative of 3 independent experiments with ≥ 95 cells analysed per condition. (**C)** $EC_{50}$ of carbachol calculated from **B**. (*** = $p$<0.001).

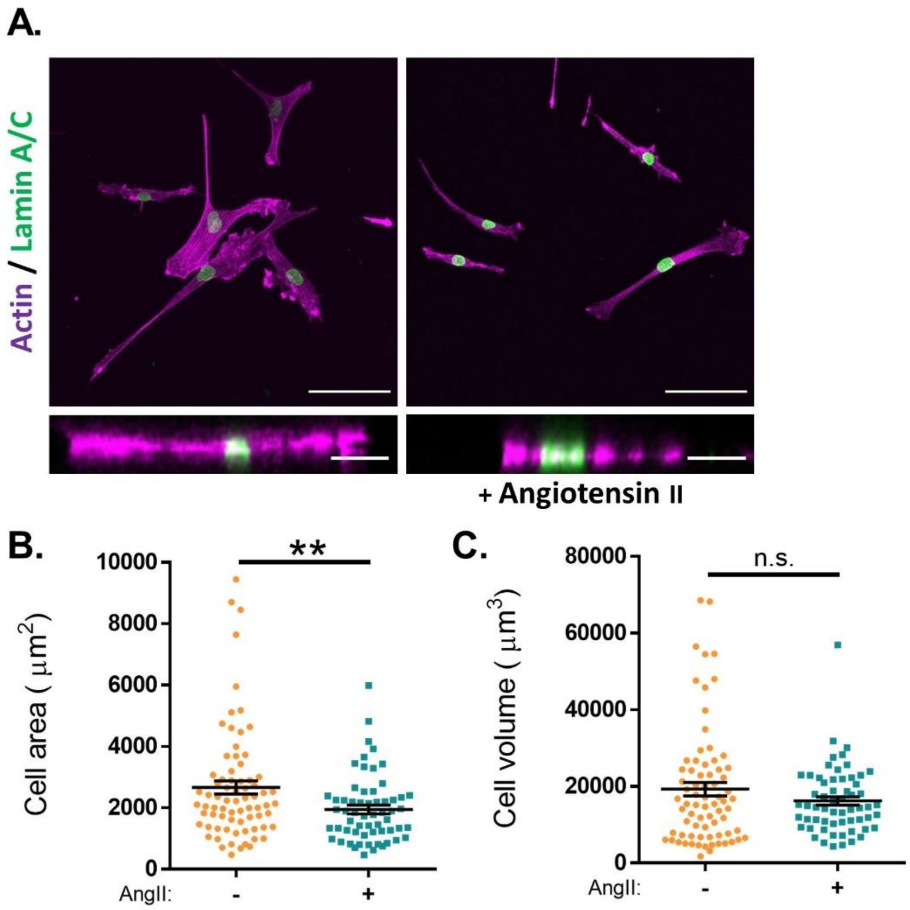


**Supplementary Figure S3: Angiotensin II stimulation promotes VSMC area to decrease but volume remains unchanged. (A)** Representative images of isolated VSMCs cultured on 12 kPa polyacrylamide hydrogels, with or without angiotensin II (AngII) stimulation for 30 minutes. Actin cytoskeleton (Rhodamine phalloidin, Purple) and Lamin A/C (green). **Top -** Representative XY images of VSMC area. Scale bar = 100 μm. **Bottom -** Representative XZ images of VSMC height. Scale bar = 30 μm. **(B)** Isolated VSMC area and **(C)** Isolated VSMC volume. Graphs are representative of 3 independent experiments with ≥ 60 cell analysed per condition. (n.s. = non-significant), (** = $p$<0.01).

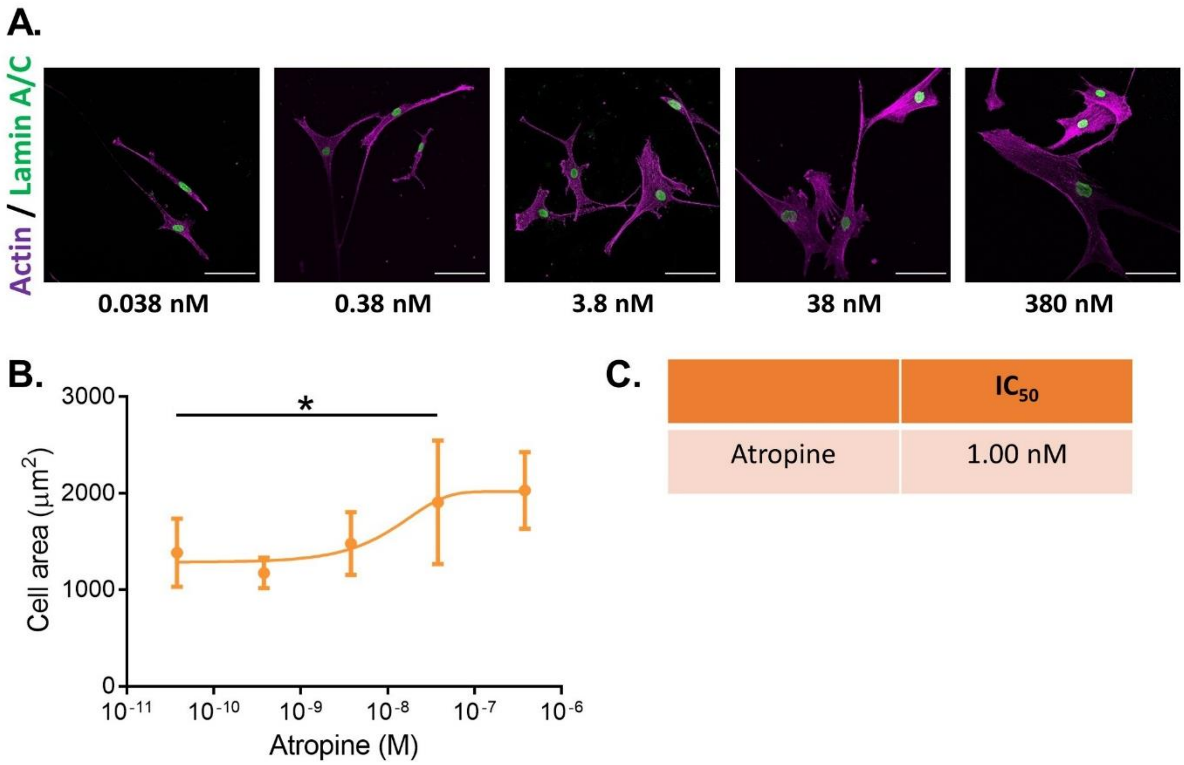


**Supplementary Figure S4: Atropine blocks carbachol mediated VSMC contraction on pliable hydrogels. (A)** Representative images of isolated VSMCs cultured on 12 kPa hydrogels and treated with carbachol (10 µM) for 30 minutes in the presence of a range of atropine concentrations. Actin cytoskeleton (Rhodamine phalloidin, Purple) and Lamin A/C (green). Scale bar = 100 µm. **(B)** Isolated VSMC area, representative of 3 independent experiments with ≥ 35 cells analysed per condition. **(C)** $IC_{50}$ of atropine calculated from **B**. (* = $p$<0.05).

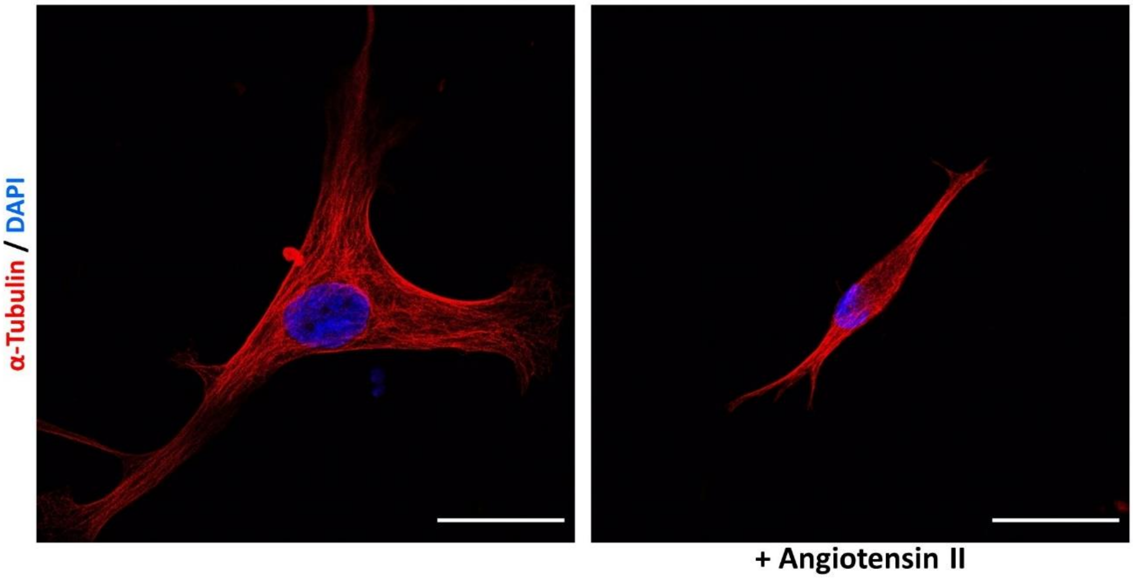


**Supplementary Figure S5: Angiotensin II stimulation induces reorganisation of the microtubule cytoskeleton.** Representative images of isolated VSMCs cultured on 12 kPa polyacrylamide hydrogels and treated with angiotensin II (10 µM) for 30 minutes. α-tubulin (Red) and cell nuclei (DAPI, Blue). Scale bar = 50 µm. Images representative of those from 3 independent experiments.

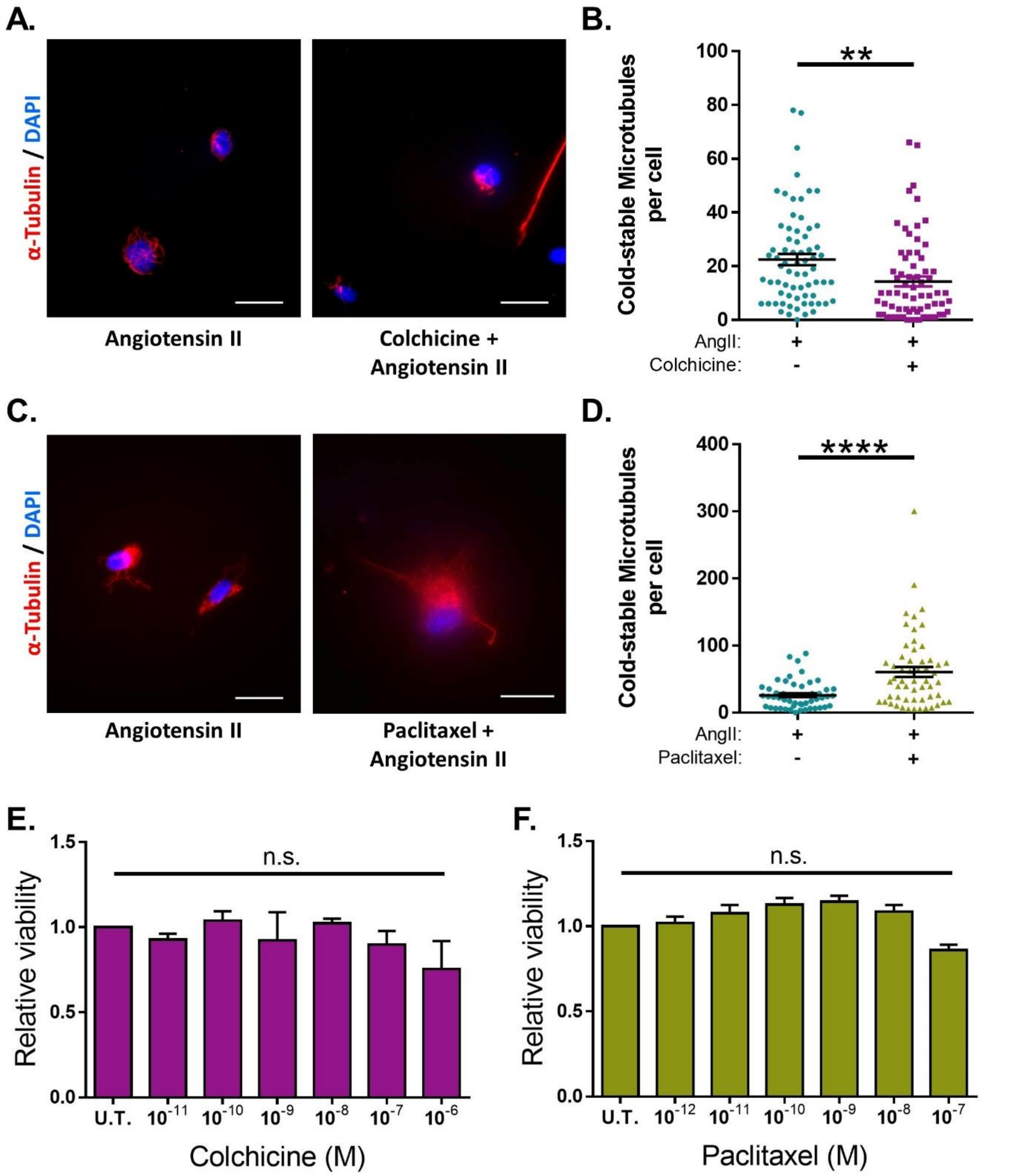


**Supplementary Figure S6: Microtubule targeting agents regulate microtubule stability in isolated VSMCs. (A)** Representative images of isolated VSMCs cultured on 12 kPa polyacrylamide hydrogels, stained for cold-stable microtubules (α-tubulin, Red) and cell nuclei (DAPI, Blue). Scale bar = 50 µm. Cells were pre-treated +/- colchicine (100 nM) for 30 minutes prior to cotreatment with angiotensin II (AngII) (10 µM) for an additional 30 minutes. **(B)** Number of cold-stable microtubules per cell, representative of 4 independent experiments, with 69 cells analysed per condition. **(C)**

Representative images of isolated VSMCs cultured on 12 kPa polyacrylamide hydrogels, stained for cold-stable microtubules (α-tubulin, Red) and cell nuclei (DAPI, Blue). Scale bar = 50 µm. Cells were pre-treated +/- paclitaxel (1 nM) for 30 minutes prior to cotreatment with angiotensin II (AngII) (10 µM) for an additional 30 minutes. **(D)** Number of cold-stable microtubules per cell, representative of 4 independent experiments, with ≥ 52 cells analysed per condition. **(E)** Relative VSMC viability following a 1 hr treatment with a range of colchicine concentrations. **(F)** Relative VSMC viability following a 1 hr treatment with a range of paclitaxel concentrations. Both **E&F** are representative of 4 independent experiments. U.T. = Untreated. (n.s. = non-significant), (** = $p$<0.01) (**** = $p$<0.0001).
